# Semaglutide-induced weight loss improves mitochondrial energy efficiency in skeletal muscle

**DOI:** 10.1101/2024.11.13.623431

**Authors:** Ran Hee Choi, Takuya Karasawa, Cesar A. Meza, J. Alan Maschek, Allison Manuel, Linda S. Nikolova, Kelsey H. Fisher-Wellmen, James E. Cox, Amandine Chaix, Katsuhiko Funai

**Author notes:** These authors contributed equally (Ran Hee Choi, Takuya Karasawa). Co-correspondence: CONTACT INFO, Katsuhiko Funai, PhD, Diabetes & Metabolism Research Center 15N, 2030E, Salt Lake City, UT 84112 University of Utah.

## Abstract

**Objective:** Glucagon-like peptide 1 receptor agonists (e.g. semaglutide) potently induce weight loss and thereby reducing obesity-related complications. However, weight regain occurs when treatment is discontinued. An increase in skeletal muscle oxidative phosphorylation (OXPHOS) efficiency upon diet-mediated weight loss has been described, which may contribute to reduced systemic energy expenditure and weight regain. We set out to determine the unknown effect of semaglutide on muscle OXPHOS efficiency.

**Methods:** C57BL/6J mice were fed a high-fat diet for 12 weeks before receiving semaglutide or vehicle for 1 or 3 weeks. The rate of ATP production and O_2_ consumption were measured by a high-resolution respirometry and fluorometry to determine OXPHOS efficiency in skeletal muscle at these 2 timepoints.

**Results:** Semaglutide treatment led to significant reductions in fat and lean mass. Semaglutide improved skeletal muscle OXPHOS efficiency, measured as ATP produced per O_2_ consumed (P/O) in permeabilized muscle fibers. Mitochondrial proteomic analysis revealed changes restricted to two proteins linked to complex III assembly (Lyrm7 and Ttc1, p <0.05 without multiple corrections) without substantial changes in the abundance of OXPHOS subunits.

**Conclusions:** These data indicate that weight loss with semaglutide treatment increases skeletal muscle mitochondrial efficiency. Future studies could test whether it contributes to weight regain.

## Introduction

Obesity predisposes individuals to a broad spectrum of diseases, including type 2 diabetes and cancer. Modest (5-10%) weight loss improves clinical parameters associated with obesity-related diseases (1). Lifestyle interventions can successfully induce ≥5% weight loss from an overweight state; however, maintaining a weight-reduced state has been extremely challenging. Almost 80% of people who have undergone weight loss by various methods is expected to undergo a weight regain phase over the next 5 years (2). Combating weight rebound for sustainable weight management remains a primary challenge, and is compounded by environmental, behavioral, and physiological factors that promote regain.

Weight loss from an overweight state increases energy efficiency thereby lowering the fuel cost during low levels of physical activity (3–5). This metabolic adaptation persists for several years following weight loss and makes individuals susceptible to significant weight regain (3, 6). It has been demonstrated that increased skeletal muscle work efficiency during as well as after weight loss contributes to decreased energy expenditure (4). Potential mechanisms underlying metabolic adaptations in the weight-reduced state include decreased glycolytic enzymatic activity, such as phosphofructokinase, lower bioactive thyroid hormones, increased activity of sarco/endoplasmic reticular Ca^2+^-dependent ATPase (SERCA), shift in myosin heavy chain (MHC) isoforms toward MHC I—characterized as more efficient and oxidative—that may enhance skeletal muscle work efficiency in a weight-reduced state by altering myofibrillar ATPases (4, 5, 7). Similar changes in metabolic rates can be found in the context of exercise training and high-altitude adaptation (8, 9).

Recently, we showed that dietary intervention-induced weight loss in overweight mice increased mitochondrial oxidative phosphorylation (OXPHOS) efficiency of skeletal muscle, quantified by ATP production per O_2_ consumption (P/O) (10). Those changes were not associated with changes in SERCA abundance or efficiency compared to lean controls and overweight groups. These results suggest that increased energy efficiency in a weight-reduced state may be due to alteration of mitochondrial OXPHOS efficiency, which requires less O_2_ to produce equimolar of ATP production.

In recent years, glucagon-like peptide-1 (GLP-1) receptor agonists, such as semaglutide, have shown striking efficacy for weight loss, inducing 10% to 15% reduction in body weight within one year. In STEP and SELECT trials lasting 68 weeks and 208 weeks respectively, weekly subcutaneous injection of semaglutide (2.4 mg) decreased body weight by approximately 10% to 15% within one year, which was successfully sustained for the experimental duration (11, 12). Importantly, individuals who stopped taking GLP-1 receptor agonists regained 10% of their lost weight within a year (13). While changes in food intake likely largely contribute to weight regain, there are limited studies on the contribution of energy efficiency/expenditure on the propensity for weight regain during or after GLP-1 receptor agonists treatment. Furthermore, neither the duration of changed energy efficiency during and/or after weight loss, as well as the outcomes of sustained increase in energy efficiency have been fully described. As mentioned above, increased energy efficiency from weight loss is like that observed with exercise training, thus increased energy efficiency in the weight-reduced state could have potential benefits on the health, such as improving glycemic controls and reducing oxidative stress (14, 15).

In this study, we set out to determine the influence of GLP-1 receptor agonist-induced weight loss on skeletal muscle mitochondrial energy efficiency. We hypothesized that semaglutide treatment will increase energy efficiency of skeletal muscle mitochondrial ATP production, similarly to our observation with dietary intervention-induced weight loss (10).

## Materials and Methods

### Animals

Male C57BL/6J (RRID:IMSR_JAX:000664) mice were purchased from Jackson Laboratory at 20 weeks of age. Mice were maintained on a high-fat diet (42% calories from fat, Inotiv TD88137) for the duration of the experiments. Body composition was determined by Bruker Minispec NMR (Bruker, Germany) before and after the interventions. At 12 weeks of HFD feeding, mice were divided into two groups: vehicle (phosphate buffered saline, PBS) and semaglutide (Novo Nordisk, s.c. 3 nmol/kg). Reagents were administered once daily for either 1-week or 3-week. Mice were housed in a 12-h light/dark cycle and provided access to food and water *ad libitum*. For terminal experiments, mice were injected intraperitoneally with ketamine (80 mg/kg) and xylazine (10 mg/kg) after which tissues were collected. All protocols were approved by Institutional Animal Care and Use Committees at the University of Utah.

### Preparation of permeabilize muscle fiber bundles (PmFB)

A small portion of mouse red gastrocnemius muscle was dissected and placed in buffer X [7.23 mM K_2_EGTA, 2.77 mM CaK_2_EGTA, 20 mM imidazole, 20 mM taurine, 5.7 mM ATP, 14.3 mM phosphocreatine, 6.56 mM MgCl_2_·6H_2_O, and 50 mM 2-(N-Morpholino) ethanesulfonic acid potassium salt (K-MES) (pH7.4)]. Fiber bundles were separated and permeabilized for 30 min at 4°C with saponin (30 µg/ml). Immediately after permeabilization, fiber bundles were washed in buffer Z [105 mM K-MES, 30 mM KCl, 10 mM K_2_HPO_4_, 5 mM MgCl_2_·6H_2_O, 0.5 mg/mL BSA, and 1 mM EGTA, pH 7.4] with 0.5 mM pyruvate and 0.2 mM malate for 15 min. After washing, fiber bundles were placed in buffer Z until analysis.

### Mitochondria isolation from mouse skeletal muscle

Mouse gastrocnemius muscle was freshly dissected and used to isolate mitochondria as previously described (16). Briefly, the tissue was minced in ice-cold mitochondrial isolation medium (MIM) buffer [0.3 M sucrose, 10 mM 4-(2-hydroxyethyl)-1-piperazineethanesulfonic acid (HEPES), 1 mM ethylene glycol tetraacetic acid (EGTA), pH 7.1] with bovine serum albumin (BSA; 1 mg/ml) and homogenized using a TH-01 homogenizer (Omni International). The homogenate was then centrifuged at 800 g for 10 min at 4°C. The supernatant was transferred to a new tube then centrifuged at 12,000 g for 10 min at 4°C. The supernatant was removed and washed the crude mitochondrial pellet in 1 ml MIM buffer. Subsequently the pellet was centrifuged again at 12,000 g for 10 min at 4°C, and the pellet was resuspended in 100 µl MIM buffer.

### High-resolution respirometry and fluorometry

PmFB and isolated mitochondria were used to determine oxygen consumption and ATP production as previously described (10). Respiration was measured using the Oroboros Oxygraph-2K (Oroboros Instruments). ATP production was determined by enzymatically coupling ATP production to NADPH synthesis using Horiba Fluorolog-QM (Horiba Scientific), as previously described (17). Both experiments were performed in buffer Z in the presence of 0.5 mM malate, 5 mM pyruvate, 5 mM glutamate, and 10 mM succinate. Respiration and ATP production was recorded with a subsequent ADP titration (20, 200, and 2000 μM for PmFB; 2, 20, and 200 μM for isolated mitochondria). In PmFB experiments, 20 mM creatine monohydrate and 10 M blebbistatin were added to buffer Z to inhibit myosin adenosine triphosphatases. The P/O ratio was analyzed by dividing the rate of ATP production by the rate of atomic O_2_ consumed (17).

### Sample preparation and nLC-MS/MS label free proteomic analysis

Mitochondria from mouse skeletal muscle were isolated and prepared for label free proteomic analysis as previously described with some modifications (10). Isolated mitochondria resuspended in MIM buffer was added an ice-cold acetone and incubated for 10 min to precipitate protein. The samples were vortexed then centrifuged at 12,000 g for 10 min at 4°C. The pellet was processed following the manufacturer’s instructions for S-Trap^TM^ (Protifi). Protein concentration was determined by BCA protein assay kit (ThermoFisher Scientific). Final peptides were resuspended in 0.1% formic acid for mass spectrometry analysis. Proteomic analysis was performed at the University of Proteomics Core.

Reversed-phase nano-LC-MS/MS was performed on a nanoElute 2 (Bruker Daltonics) coupled to a Bruker timsTOF Pro2 mass spectrometer equipped with a nanoelectrospray source. 150 ng of each sample was injected directly onto the liquid chromatograph reverse-phase C18 15 cm x 150 µm PepSep^TM^ nanocolumn (Bruker Daltonics) heated to 50°C. The peptides were eluted with a gradient of reversed-phase buffers (Buffer A: 0.1% formic acid in 100% water; Buffer B: 0.1% formic acid in 100% acetonitrile) at a flow rate of 0.5 µL/min. The LC run lasted for 45 minutes with a starting concentration of 5% buffer B increasing to 28% buffer B over 40 minutes, up to 32% buffer B over 2 minutes and held at 95% B for 3 minutes. The separation column is equilibrated with 4 column volumes at 800 bar following each run. The timsTOF Pro2 was operated in PASEF data-dependent acquisition (DDA) MS/MS scan mode to generate the custom peptide library. The TIMS section was operated with a 120 ms ramp time at a rate of 7.93 Hz and an ion mobility scan range of 0.6-1.4 V·s/cm^2^. MS and MS/MS spectra were recorded from 100 to 1,700 m/z. A polygon filter was applied to select against singly charged ions. The quadrupole isolation width was set to 3 Da. The mass spectrometer was operated in PASEF data-independent acquisition (DIA) MS/MS scan mode to analyze each experimental sample. The TIMS section was operated with a 120 ms ramp time at a rate of 7.93 Hz and an ion mobility scan range of 0.6-1.4 V·s/cm^2^. dia-PASEF window parameters were set to mass width of 25 Da and 47 mass steps/ cycle. MS and MS/MS spectra were recorded from 147 to 1,322.6 m/z. The custom peptide library was created using Fragpipe v20.0 software (downloaded 4-7-23) against the uniprot_ref_mouse_database (downloaded 12-7-2023 with 17,191 proteins) as well as MitoCarta 3.0 (Rath 2021). Protein abundances based on peak intensity were calculated using DIA-NN software version 1.8.1. An allowance was made for 1 missed cleavage following trypsin/Lys C digestion. No fixed modifications were considered. The variable modifications of methionine oxidation, N-terminal acetylation and cysteine carbamidomethylation (IAA) were considered with a mass tolerance of 15 ppm for precursor ions and a mass tolerance of 10 ppm for fragment ions. The results were filtered with a false discovery rate of 0.01 for both proteins and peptides. Mitochondrial enrichment factor was determined by comparing mitochondrial protein abundance to total protein abundance.

### Mitochondrial lipid mass spectrometry

Mitochondrial lipidome was analyzed from isolated mitochondria from mouse gastrocnemius muscle. Lipids were extracted as described previously (10). Briefly, a mixture of ice-cold methyl-tert-butyl ether (MTBE), methanol, and internal standards (EquiSPLASH, Avanti, Cat #: 330731 and Cardiolipin Mix I, Avanti, Cat #: LM6003) was added to 50 µg of mitochondrial protein. The samples were vortexed and sonicated for 1 min. The samples were then incubated on ice for 15 min and vortexed every 5 min. Phase separation was induced by adding H_2_O to the samples, then the 15-min incubation with vortex every 5 min was performed. The samples were centrifuged at 12,000 g for 10 min. The organic phase was collected to a new tube while the aqueous phase was used for second extraction with 1 ml of 10:3:2.5 (v/v/v) MTBE/methanol/H_2_O. The organic layers were combined and dried under Genevac^TM^ miVac centrifugal evaporation system. The dried lipids were reconstituted in 300 ul of 8:2:2 (v/v/v) IPA/ACN/H_2_O. Mitochondrial lipidome profiles were obtained using targeted LC-MS/MS platforms in operation at the University of Utah Metabolomics Core. Lipids contents were normalized to mitochondrial protein levels.

### Western blot

Proteins from isolated mitochondria and whole frozen muscle were utilized for the analysis. Frozen gastrocnemius muscle was homogenized using TH-01 homogenizer in ice-cold RIPA buffer (ThermoFisher Scientific, Cat #: 89901) supplemented with protease inhibitor cocktail (ThermoFisher Scientific, Cat #: 78446). Protein concentration was determined by BCA protein kit (ThermoFisher Scientific). Equal amounts of proteins were analyzed with 4-20% gradient SDS-polyacrylamide gel (Bio-Rad) for electrophoresis and transferred to polyvinylidene fluoride membranes (ThermoFisher Scientific). The membranes were blocked with 3% skim milk in Tris-buffered saline containing 0.1% Tween-20 (TBST) for 1 hr at room temperature, followed by incubation of primary antibodies, OXPHOS (Abcam, MS604-300), at 4°C overnight. After overnight, the membranes were washed with TBST and incubated with secondary antibody (1:5,000) in 1% skim milk for 1 hr at room temperature. Following three washes in TBST and one in TBS, the membranes were incubated with Western Lightning Plus-ECL (PerkinElmer) and detected by a ChemiDoc Imaging System (Bio-Rad) and quantified with Image Lab Software (Bio-Rad).

### Native PAGE

Mitochondria supercomplex analysis was performed as previously described with modification (18). Isolated mitochondria (25 μg) suspended in MIM were pelleted at 12,000 x g for 10 min and subsequently solubilized in 20 μL sample buffer (5 μL of 4x Native PAGE Sample Buffer, 2 μL 10% digitonin, 13 μL ddH_2_O per sample) for 20 min on ice and then centrifuged at 18,000 x g for 10 min at 4°C. The supernatant (15 μL) was collected and placed into a new tube and mixed with 1 μL of G-250 sample additive (Thermo Fisher Scientific). The samples and protein ladder were loaded onto a native PAGE 3-12% Bis-Tris Gel (Thermo Fisher Scientific). Dark blue cathode buffer (0.02% Coomassie Brilliant Blue dissolved in Native Page running buffer) was carefully added to the front of gel box and Native PAGE running buffer (anode buffer) was carefully added to the back of the gel box making sure to not mix. Electrophoresis was performed at 150 V for 45 min at 4°C. The dark blue cathode buffer was then replaced with light blue cathode buffer (Dark blue cathode buffer and anode buffer 1:9) and run at 15 mA for 90 min at 4°C. Gels were subsequently washed with transfer buffer (50 mM tricine, 7.5 mM imidazole) for 30 min and transferred to PVDF membrane at 100 V for 60 min. The membranes were stained with Coomassie Brilliant Blue staining solution (0.05% Coomassie Brilliant Blue, 50% methanol, 10% acetic acid) and washed destaining solution (50% methanol, 7% acetic acid) to visualize the membrane bound protein and ladder. The detection of OXPHOS protein was performed with the same procedure described in the Western blot section.

### Electron microscopy

Samples preparation for electron microscopy was performed as previously described (16). Briefly, freshly dissected red gastrocnemius muscles from 3 weeks of vehicle and semaglutide were placed in fixative solution (1% glutaraldehyde, 2.5% paraformaldehyde, 100 mM cacodylate buffer pH 7.4, 6 mM CaCl_2_, 4.8% sucrose) and cut into sections ∼2 mm then stored at 4°C for 48 hours. Samples were prepared by the Electron Microscopy Core at University of Utah. The tissues underwent 3 × 10-minute washes in 100 mM cacodylate buffer (pH 7.4) prior to secondary fixation (2% osmium tetroxide) for 1 hour at room temperature. Osmium tetroxide as a secondary fixative has the advantage of preserving membrane lipids, which are not preserved using aldehyde, alone. After secondary fixation, samples were subsequently rinsed for 5 minutes in cacodylate buffer and distilled H_2_O, followed by prestaining with saturated uranyl acetate for 1 hour at room temperature. After prestaining, each sample was dehydrated with a graded ethanol series (2 × 15 minutes each: 30%, 50%, 70%, 95%; then 3 × 20 minutes each: 100%) and acetone (3 × 15 minutes) and were infiltrated with EPON epoxy resin (5 hours 30%, overnight 70%, 3 × 2-hour 40 minute 100%, 100% fresh for embed). Samples were then polymerized for 48 hours at 60°C. Ultracut was performed using Leica UC 6 ultratome with sections at 70 nm thickness and mounted on 200 mesh copper grids. The grids with the sections were stained for 20 minutes with saturated uranyl acetate and subsequently stained for 10 minutes with lead citrate. The sections were imaged at 120 kV using JEOL-JEM 1400plus (Tokyo, Japan) transmission electron microscope and images were acquired on a Gatan 2kx2k digital camera (Gatan, Inc.)

### Statistical analysis

Statistical analysis was performed using GraphPad Prism 10.2.3 software. Two-way ANOVA followed by Tukey’s multiple comparisons test or unpaired t-test were performed for group comparison. All data are presented as mean±SEM, and statistical significance was set at p ≤0.05.

## Results

### Semaglutide treatment induces weight loss in overweight mice

Wildtype C57BL/6J mice were fed a high-fat-diet (HFD, 42% kcal from fat) for 12 weeks before being treated with either vehicle (PBS) or semaglutide (3 nmol/kg body weight/day, subcutaneous injection) for 1 week or 3 weeks (Fig. 1A). During the intervention, mice were maintained on the HFD. Mice treated with semaglutide lost about 12% ± 1.7 of body weight after 1 week of treatment and about 23 % ± 1.66 after 3 weeks (Fig. 1B and 1C). Loss in body weight was due to disproportionately more loss in fat mass than lean mass. Fat mass was significantly decreased by 26% ± 2.23 after 1 week and by 42% ± 3.66 after 3 weeks of semaglutide treatment compared to the vehicle groups (Fig. 1D and 1E). Lean mass was significantly reduced by 9% ± 1.5 both after 1 week and 3 weeks (Fig. 1F and 1G) and this similarly reflected in gastrocnemius muscle mass (Fig. 1H). Thus, semaglutide intervention significantly reduced body weight, fat mass, and lean mass.

**Figure 1.**
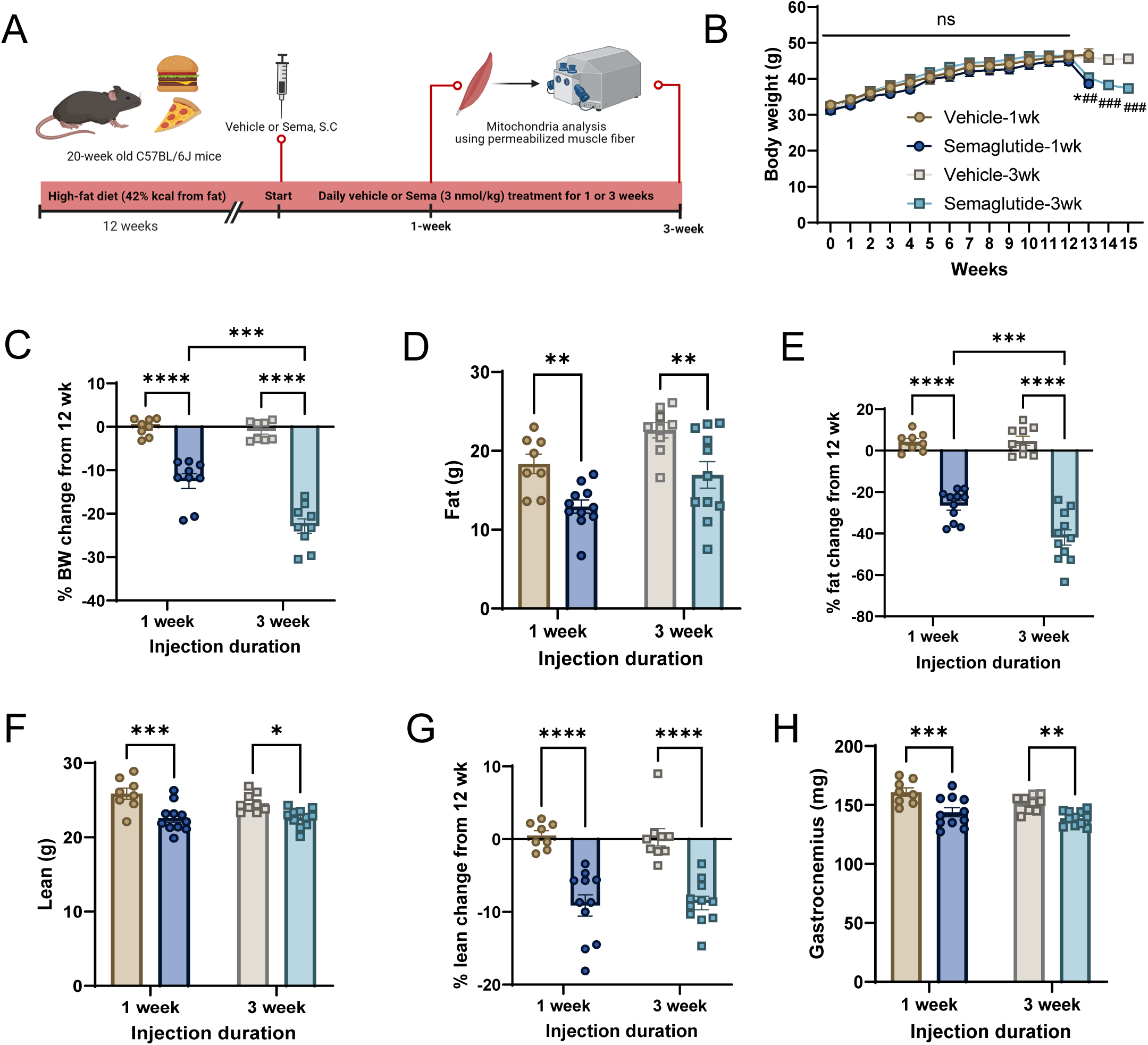
Semaglutide induces weight loss in overweight mice. (A) Schematic of the experimental design. Twenty-week-old C57BL/6J mice were fed 42% HFD diet for 12 weeks before semaglutide (3 nmol/kg of body weight/day) intervention. Semaglutide was subcutaneously injected daily for 1 or 3 weeks. Vehicle groups received an equal volume of PBS. (B) Body weight was recorded every week. (C) Percent change in body weight after semaglutide treatment was calculated from the baseline of 12-wk HFD. (D) Fat mass was determined by NMR before and after treatment. (E) Percentage change in fat mass was analyzed by NMR. (F) Lean mass was determined by NMR before and after treatment. (G) Percentage change in lean mass was calculated by NMR. (H) After the completion of semaglutide treatment, gastrocnemius muscles were harvested, and wet weight was measured. Data are represented as mean± SEM. * p<0.05, ** p<0.01, *** p<0.001, **** p<0.0001 significant difference vs. corresponding controls. ## p<0.01, ### p<0.001 significant difference vs. vehicle-3wk. N=8 and 9 for vehicle-1wk and vehicle-3wk, respectively, and N=11 for both semaglutide 1-wk and 3-wk.

### Semaglutide-induced weight loss increases skeletal muscle OXPHOS energy efficiency

To test whether semaglutide-induced weight loss increases the energy efficiency for mitochondrial ATP synthesis, we first quantified the rates of ATP production, O_2_ consumption, and P/O in permeabilized red gastrocnemius muscle fiber bundles harvested from 1-week and 3-week vehicle and semaglutide-treated groups. The rate of ATP production (*J*ATP) showed a trend toward an increase after 1 week of semaglutide treatment, with a more pronounced effect observed after 3 weeks of treatment (Fig. 2A and 2B). Additionally, while O_2_ consumption (*J*O_2_) did not change significantly after 1 week, it was markedly decreased after 3 weeks of semaglutide treatment (Fig. 2C and 2D). When calculating the P/O ratio to determine energy efficiency, semaglutide tended to increased P/O after 1 week, with a significant improvement after 3 weeks (Fig. 2E and 2F). This improved P/O was not associated with changes in OXPHOS subunits levels (Fig. 2G and 2H). Overall, these results indicate that 3 weeks of semaglutide treatment robustly improved mitochondrial OXPHOS efficiency in skeletal muscle by reducing *J*O_2_ without affecting OXPHOS subunits.

**Figure 2.**
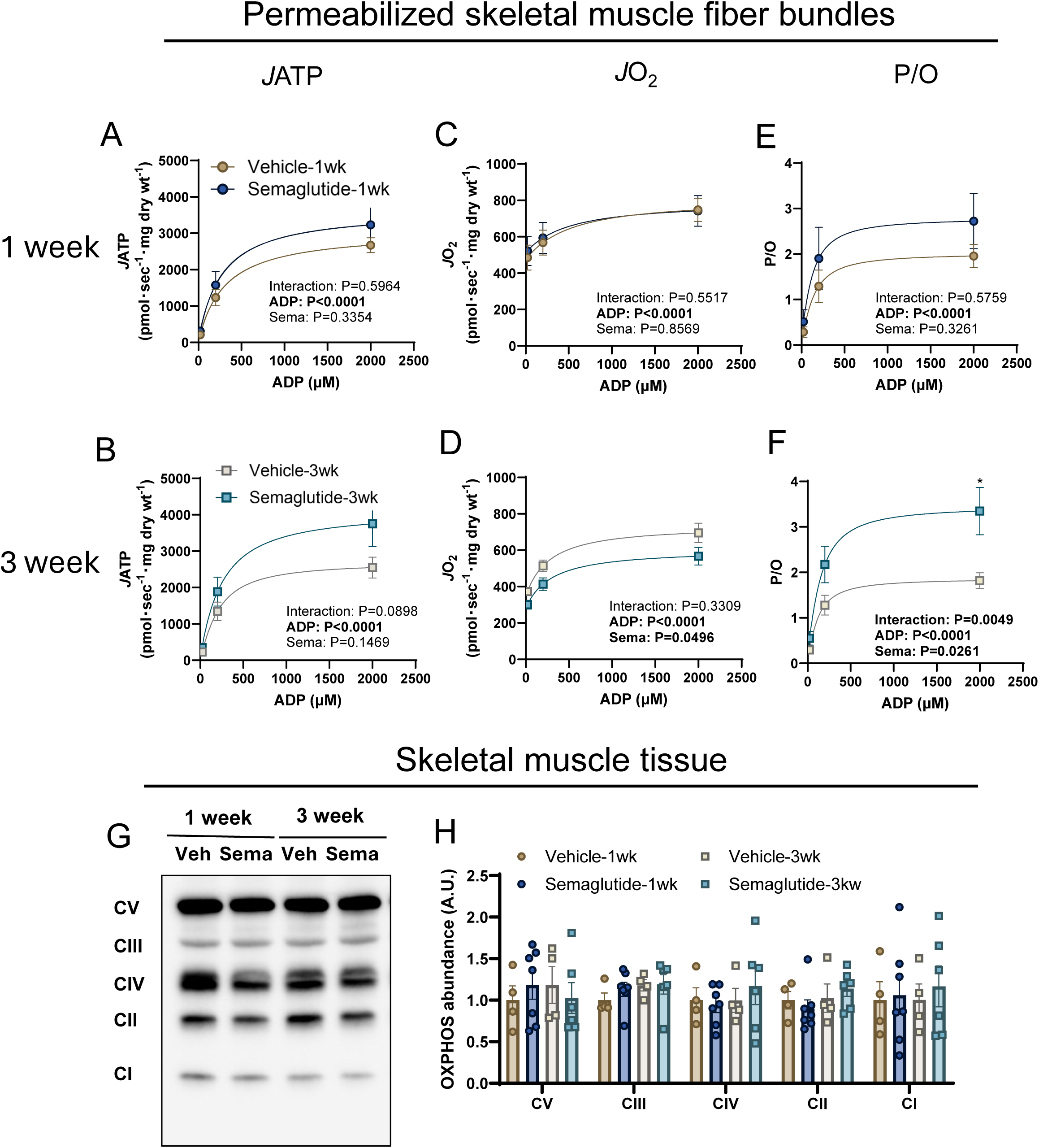
Semaglutide treatment increases skeletal muscle OXPHOS energy efficiency. A small piece of red gastrocnemius was collected and permeabilized in the buffer with saponin (30 µg/ml). (A and B) The rate of ATP production (*J*ATP) was determined by fluorometer. (C and D) Mitochondrial O_2_ consumption (*J*O_2_) was analyzed by high-resolution respirometry. (E and F) OXPHOS efficiency was determined as P/O ratio. (G and H) Abundance of OXPHOS subunits in whole lysate of gastrocnemius muscle was determined by Western blot. Data are represented as mean± SEM. * p<0.05 significant difference vs. corresponding controls. N=10 and 13 for vehicle-1wk and vehicle-3wk and N=10 and 14 for semaglutide 1-wk and 3-wk, respectively.

### Semaglutide-induced increase in mitochondrial OXPHOS efficiency is NOT observed in isolated mitochondria from skeletal muscle

Mitochondrial bioenergetic phenotyping in permeabilized muscle fiber bundles enables assessment of respiratory responses in situ, better preserving the native conformation of mitochondrial reticulum and membrane structures. However, these preparations also contain other subcellular structures, including other ATP synthases and ATPases. While respiratory assay buffers contain inhibitors for some of these systems, we cannot rule out the possibility that non-mitochondrial mechanisms could influence ATP fluorometry (less concerning for O_2_ consumption) and thereby affect P/O measurements.

To address this, we conducted the same bioenergetic phenotyping in isolated mitochondria from gastrocnemius muscles. In isolated mitochondria, neither *J*ATP, *J*O_2_, nor P/O were significantly changed by semaglutide (Fig. 3A-F), and OXPHOS subunits remained unchanged (Fig. 3G and 3H). We observed that *J*ATP responses stopped after the second dose of ADP (20 μM) (Fig. 3A and 3B), unlike what was observed in permeabilized muscle fibers (Fig. 2A and 2B), while *J*O_2_ continued to respond to each ADP titration (Fig. 3C and 3D). This led to a decrease in P/O at the highest ADP compared to the second dose (Fig. 3E and 3F). These results suggest that cellular compartments associated with mitochondrial network preserved in permeabilized muscle fiber bundles may play a critical role in the improvement of semaglutide-induced OXPHOS efficiency in skeletal muscle.

**Figure 3.**
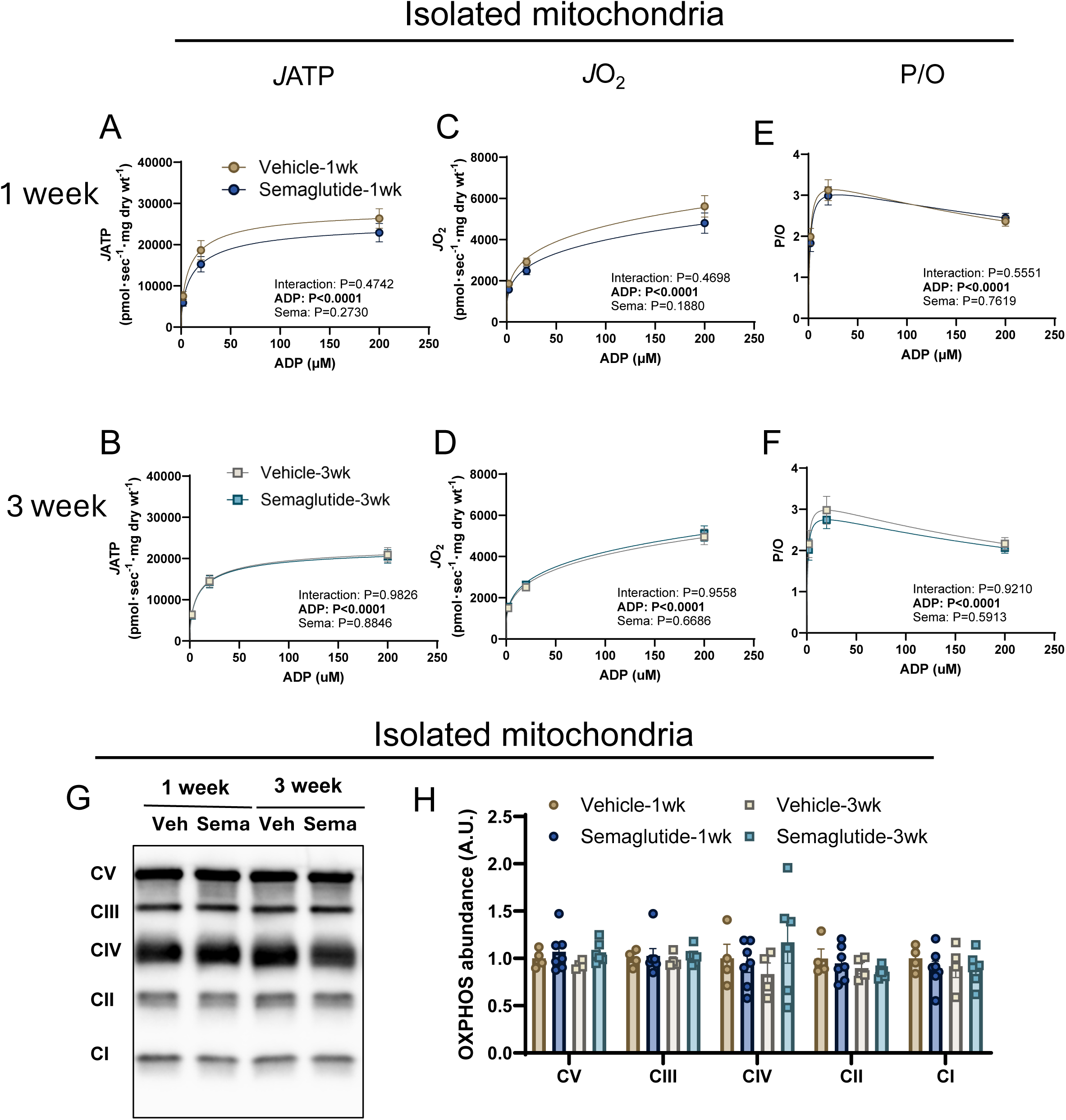
Semaglutide-induced increase in mitochondrial OXPHOS efficiency is NOT observed when quantified in isolated mitochondria from skeletal muscle. Mitochondria were isolated from whole gastrocnemius muscle. (A and B) The rate of ATP production (*J*ATP) was determined by fluorometer. (C and D) Mitochondrial O_2_ (*J*O_2_) consumption was analyzed by high-resolution respirometry. (E and F) OXPHOS efficiency was determined as P/O ratio. (G and H) Abundance of OXPHOS subunits in isolated mitochondria from gastrocnemius muscle was determined by Western blot. Data are represented as mean± SEM. N=12 and 13 for vehicle-1wk and vehicle-3wk, respectively, and N=15 for both semaglutide 1-wk and 3-wk.

### Semaglutide treatment does not alter skeletal muscle mitochondrial supercomplexes assembly

Mitochondrial respiratory complexes are thought to have supercomplexes that have improved efficiency for electron transport compared to free complexes (19, 20). However, semaglutide treatment did not appear to influence mitochondrial OXPHOS supercomplex assembly (Fig. 4A-F and S2A-F). Mitochondrial morphology, including shape and size of individual mitochondrion as well as those of cristae have a strong influence on OXPHOS (21). Mitochondrial morphology analyzed by electron microscopy (EM) however did not show any noticeable differences between vehicle and semaglutide treatment (Fig. 4G). Previously we have shown that mitochondrial lipidome can also influence OXPHOS (10, 16, 22), but semaglutide did not influence the mitochondrial lipidome (Fig. 5A and S1A-I). These findings indicate that alterations in mitochondrial supercomplexes formation, morphology, and lipidome are unlikely to be responsible for the improvement in muscle P/O induced by semaglutide.

**Figure 4.**
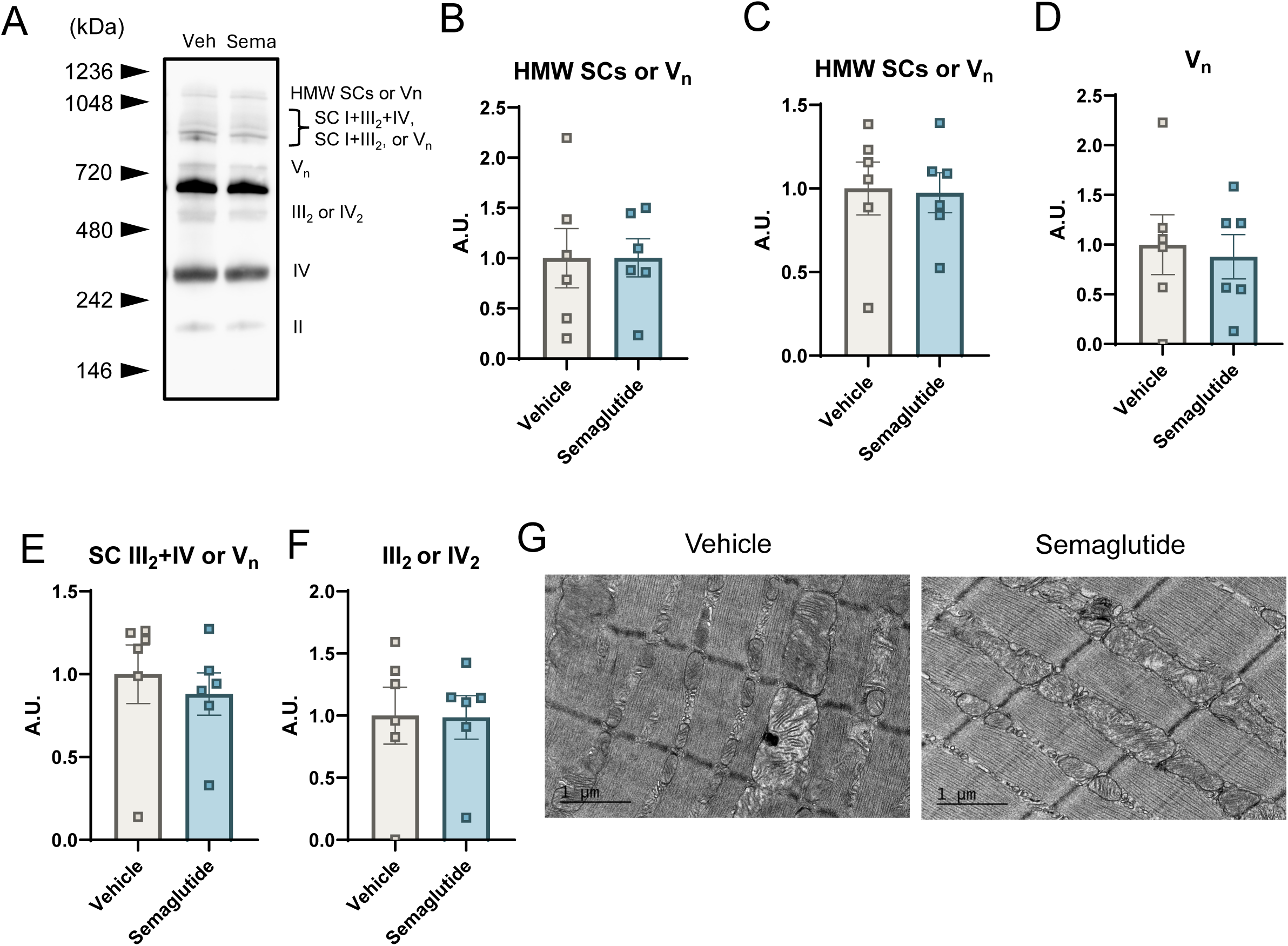
Semaglutide treatment does not alter skeletal muscle mitochondrial supercomplexes formation. (A-F) Mitochondrial supercomplexes formations were analyzed by Native PAGE and quantified. (G) Representative mitochondrial morphology analyzed by electron microscopy. All data are from 3-wk semaglutide treatment. N=6 for both vehicle and semaglutide for Native PAGE.

**Figure 5.**
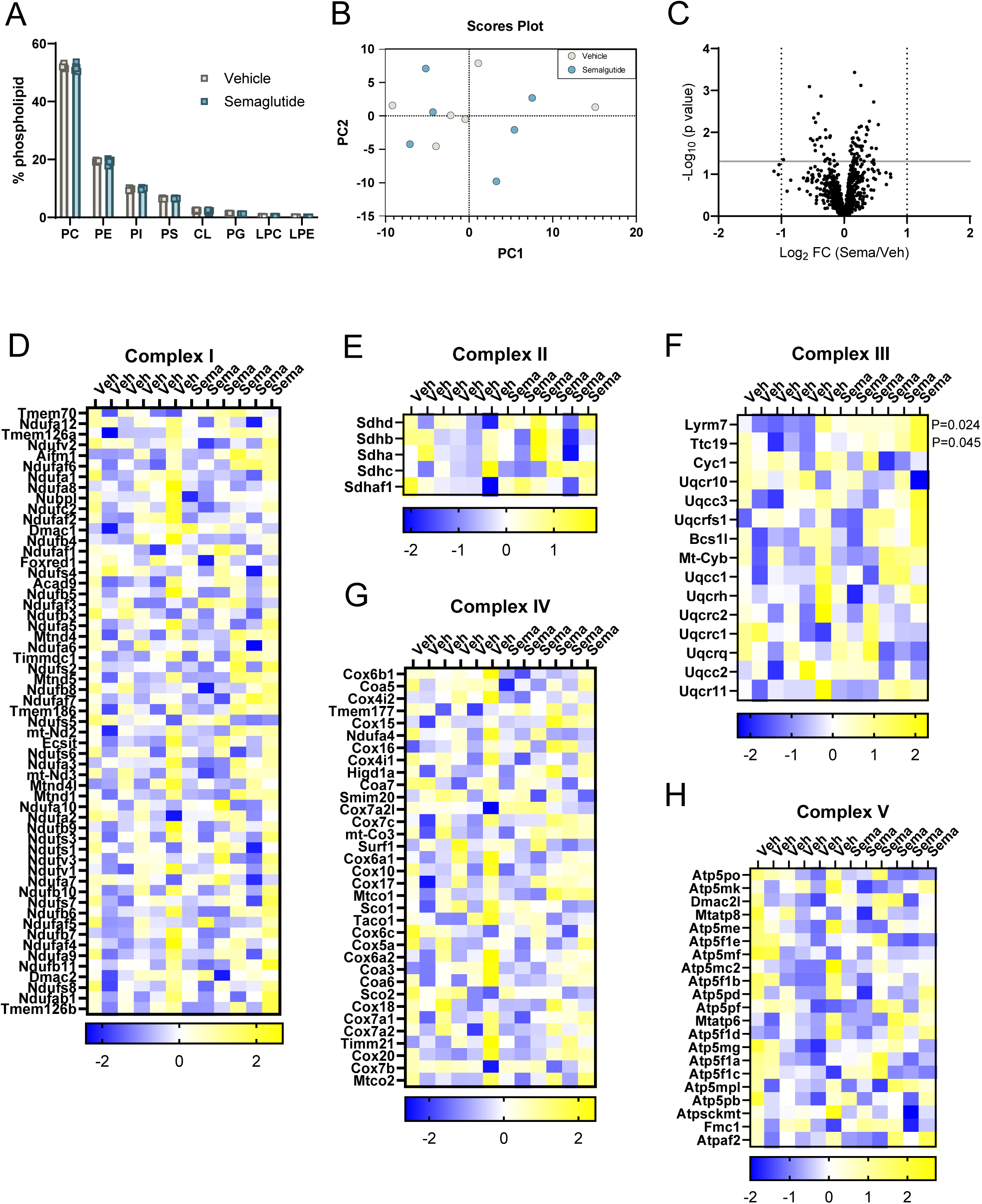
Semaglutide-induced weight loss does not robustly influence the abundances of OXPHOS subunits. (A) Relative abundance of mitochondrial lipids. (B) Principal component analysis of mitochondrial proteomic data. (C) Volcano plot of differentially abundant mitochondria proteins between vehicle (Veh) and semaglutide (Sema) groups. (D-H) Heatmap of abundance of OXPHOS subunits determined by mass spectrometry. All data are from 3-wk semaglutide treatment. N=6 for both vehicle and semaglutide.

### Semaglutide-induced weight loss does not robustly influence the abundances of OXPHOS subunits

The improvement in OXPHOS efficiency may result from changes in the abundance of respiratory subunits. Thus, we performed proteomic analyses on skeletal muscle mitochondria with or without semaglutide treatment. Consistent with our previous experiments with dietary intervention-induced weight loss (10), semaglutide treatment neither appeared to influence muscle mitochondrial proteome by principal component analyses (Fig. 5B) nor were there a single mitochondrial protein whose abundance was statistically significantly different (q<0.05) (Fig. 5C-H). However, two subunits associated with complex III assembly were trending to be elevated (Lyrm7: increased by 41%, Ttc19: increased by 38%, both p<0.05 without correcting for multiple comparison) (Fig. 5F). It is unclear whether changes in these complex III subunits are sufficient to explain the improvement in skeletal muscle OXPHOS efficiency with semaglutide treatment.

## Discussion

GLP-1 receptor agonists including semaglutide have revolutionized obesity-related health care by its efficacy to induce weight loss, as well as other comorbidities including diabetes, cardiovascular disease, and chronic kidney disease. Weight loss, including those induced by dietary intervention or those induced by GLP-1 receptor agonists, are known to induce a reduction in energy expenditure. The consequence of a perpetual reduction in resting and exercise-induced energy expenditure extends beyond the propensity for weight regain, as they may attenuate all secondary benefits of elevated energy expenditure such as glycemic control. With the emergence of GLP-1 receptor agonists and other weight loss drugs, it is important more than it has ever been to study the biology of weight-reduce state. In this study, we showed that semaglutide promotes an increase in skeletal muscle OXPHOS energy efficiency, in a similar fashion to our previous observation with weight loss induced by a dietary intervention.

The mechanism by which semaglutide treatment improved OXPHOS efficiency remains to be fully elucidated, but we found two components of complex III, Lyrm7 and Ttc19, to become elevated after semaglutide treatment.

We found that semaglutide treatment improved skeletal muscle mitochondrial efficiency when analyzed in permeabilized muscle fiber bundles. *J*ATP tended to increase after 1 week of treatment without a change in *J*O_2_, and this effect became more pronounced after 3 weeks, with significantly attenuated *J*O_2_. These changes resulted in approximately a 50% increase in the P/O ratio, indicating improved OXPHOS efficiency. These findings align with previous studies showing that weight loss, whether induced by diet or exercise, can improve mitochondrial efficiency in skeletal muscle (10, 23). Consistent with our previous study, the higher P/O ratio was driven by a reduction in *J*O_2_ in response to the same ADP stimuli (10). We also observed a trend toward increased *J*ATP with semaglutide treatment. Activation of GLP-1 receptor signaling enhances ATP synthesis in pancreatic β-cells (24, 25). Nevertheless, increased *J*ATP was not observed in our previous paper with dietary-intervention induced weight loss (10). Our previous dietary-intervention study induced weight loss by switching HFD back to standard chow, while the mice remained on HFD for our current study with semaglutide-induced weight loss. Thus, these nuanced differences between dietary-intervention and semaglutide treatment may be due to differences in their dietary compositions.

No significant improvements in mitochondrial efficiency were detected when measuring OXPHOS efficiency in isolated mitochondria. This may suggest the critical role of the intact cellular environment, which is preserved in permeabilized muscle fibers but disrupted during mitochondrial isolation (26, 27). Permeabilized muscle fiber bundles (PmFB) maintain the integrity of key cellular components, including mitochondrial structural arrangement, intracellular organelles, and multi-enzyme systems essential for energy transfer (26, 28, 29). Intact myofibrillar ATPases, including myosin ATPase, in PmFB potentially lead to this difference. We used blebbistatin, which inhibits specifically myosin II ATPase, to prevent spontaneous contraction during the measurement (17, 30). There is a possibility that the addition of blebbistatin did not inhibit other myosin ATPases, such as myosin I ATPases, presented in PmFB since we precisely dissected red gastrocnemius (type I; oxidative) for PmFB. Moreover, it has been reported that increased skeletal muscle work efficiency during, as well as after, weight loss has been linked to switch myosin isoforms from more glycolytic (IIa, IIb and IIx) to oxidative (I) (4). Other candidates for intact myofibrillar ATPases in PmFB, including Ca^2+^-activated myosin ATPase and SERCA, can also involve in generating this different observation between PmFB and isolated mitochondria as they are largely participated in ATP consumption. To avoid this possibility, we added a chelating agent to our experimental buffer (see details in Methodology). Different types of fiber or myofibrillar ATPases possibly result in distinct outcomes (27, 31, 32). This likely explains why observed improvements were specific to PmFB underscoring the importance of studying mitochondrial function in a more physiological relevant context. These findings suggest the future studies should investigate how different fiber types and mitochondria sources contribute to OXPHOS efficiency under weight loss conditions.

Semaglutide-induced improvement in OXPHOS efficiency did not appear to be related to changes in the mitochondrial lipidome. Our previous study found shifts in some cardiolipin species after diet-induced weight loss to potentially mediate increased OXPHOS efficiency (10). However, we did not observe any changes in mitochondrial phospholipids with semaglutide intervention. It remains possible that semaglutide influences other aspects of skeletal muscle mitochondria to influence bioenergetics. One possibility is that GLP-1 receptor agonists decrease mitochondrial electron leak. In leukocytes from patients with type 2 diabetes, GLP-1 receptor agonists reduced the reactive oxygen species production (33), potentially suggesting that it might reduce mitochondrial electron leak to improve efficiency. However, electron leak usually accounts for a very small portion of total electron flux (often less than 1% when quantified) (34). Thus, it is difficult to imagine that a 2-fold increase in P/O is completely accounted for by reduced electron leak. GLP-1 receptor agonists are also known to affect mitochondrial dynamics in substantia nigra (35). Even though we did not observe a robust change in mitochondrial morphology, there may be other subtle effects not detected by EM that contribute to improved OXPHOS efficiency.

Semaglutide treatment resulted in a 12% and 23% reduction in body weight after 1 and 3 weeks of treatment, respectively. Fat mass was reduced by 26% and 42% reduction over the same time point. Additionally, lean mass decreased by approximately 10% at both time points. These findings are consistent with previous clinical trials and animal studies showing that GLP-1 receptor agonists effectively promote weight loss (11, 36). Skeletal muscle is a significant contributor to energy expenditure (3, 37). A decrease in skeletal muscle mass during weight loss has been associated with a reduction in energy expenditure (3). In addition to this effect, energy expenditure per unit of lean mass appears to be reduced, promoted by an increase in skeletal muscle work efficiency (4, 5, 10).

In the current study, semaglutide did not significantly alter the skeletal muscle mitochondrial proteome when corrected for multiple comparisons. However, we identified two proteins related to complex III assembly—Lyrm7 and Ttc19—that trended to be greater (p<0.05 with t-tests). Lrym7 is involved in the incorporation of the iron-sulfur cluster into Rieske (Fe-S) protein (UQSRFC1) (38). Ttc19 is known to interact with UQCRC1 and UQCRFS1 (39), though semaglutide did not influence the abundance of these subunits. Defects in Lyrm7 and Ttc19 are linked to complex III deficiencies, nuclear type 8 (MC3DN8) and nuclear type 2 (MC3DN2), respectively, which result in increased oxidative stress and inefficient electron transfer (40–42). The association between weight loss and these proteins is unknown. Thus, semaglutide may influence Lyrm7 and Ttc19, potentially influencing complex III to enhance electron transport efficiency and reducing oxidative stress.

In summary, our study demonstrates that semaglutide treatment significantly improves mitochondrial OXPHOS efficiency in skeletal muscle. Increased OXPHOS efficiency was only observed when quantified in permeabilized fibers, suggesting that intact subcellular structures may be necessary for this effect. In contrast to our previous findings with dietary intervention-induced weight loss, we did not find semaglutide to influence skeletal muscle mitochondrial proteome or lipidome. Alternatively, improved mitochondrial energy efficiency may be mediated by post-translational modifications or by changes in mitochondria dynamics and ultrastructure. While tools to study these changes in vivo are currently limited, future studies should explore alternative molecular mechanisms that drive the improved OXPHOS efficiency with semaglutide treatment. Greater understanding of these effects could help identify targets to mitigate risks for weight regain.

## DISCLOSURE

The authors declared no conflict of interest.

## AUTHOR CONTRIBUTIONS

R.H.C, T.K, A.C, and K.F. designed the study. R.H.C, T.K, C.A.M, K.H.-F-W performed the mouse and mitochondrial experiments, and analyzed data. J.A.M. and J.E.C conducted lipidomics analyses. A.M, J.E.C, K.H.F-W conducted proteomic analysis. L.S.N assisted with electron microscopic images. R.H.C and K.F wrote the manuscript with edits from all authors.

### What is already known about this subject?

– GLP-1 receptor agonists successfully reduce adiposity in mice and in humans.
– In rodents, dietary intervention-induced weight loss improves skeletal muscle OXPHOS energy efficiency.

### What are the new findings in your manuscript?

– Semaglutide-induced weight loss also increased muscle mitochondrial OXPHOS efficiency in mice.
– Semaglutide-induced increase in mitochondrial efficiency was only observed in permeabilized muscle fibers but not in isolated mitochondria.

### How might your results change the direction of research or the focus of clinical practice?

– GLP-1 receptor agonists likely promote reduced energy expenditure in a similar fashion to dietary intervention-induced weight loss. This may influence their propensity for weight regain or other beneficial effects of increasing energy expenditure.

## Acknowledgement

This research is supported by NIH grants (DK107397, DK127979, GM144613, and AG074535 to K.F., CA278826 to K.H.F, CA286584 and AG065993 to A.H.C), and the Grant-in-aid for Japan Society for Promotion of Science (JSPS) Fellows (24KJ2039 to T.K.). University of Utah Metabolomics Core Facility is supported by DK110858.

## Figure Legends

**Figure S1.**
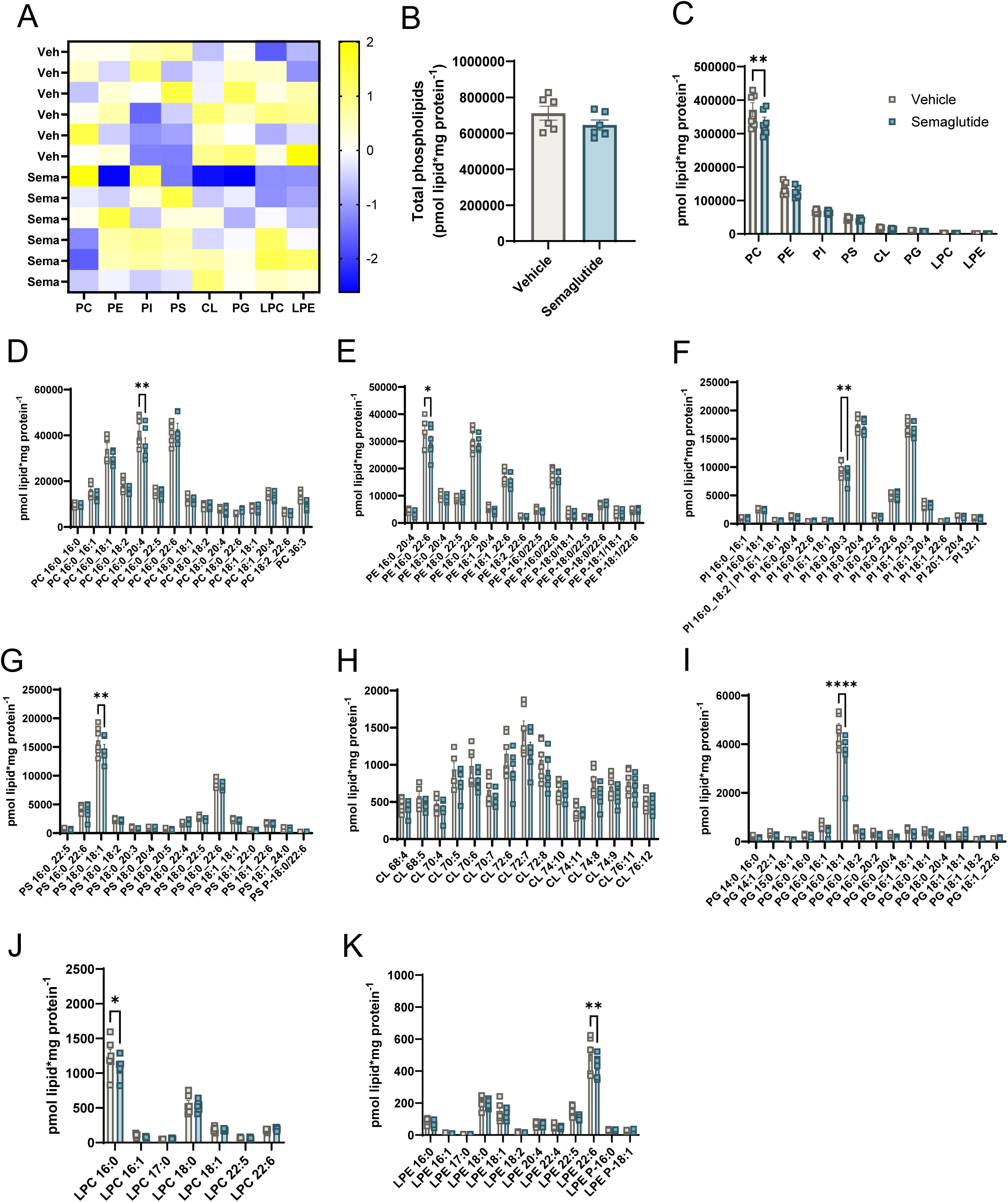
Skeletal muscle mitochondrial lipidome. Mitochondrial lipidome was analyzed in isolated mitochondrial from gastrocnemius muscle of 3 weeks vehicle and semaglutide groups. (A) Heatmap of abundance of each mitochondrial phospholipidome (B) Total phospholipids from vehicle and semaglutide group (C) Absolute abundance of mitochondrial lipids (D – K) Absolute abundance of individual phospholipids species (D) Phosphocholine (PC), (E) Phosphatidylethanolamine (PE), (F) Phosphatidylinositol (PI), (G) Phosphatidylserine (PS), (H) Cardiolipin (CL), (I) Phosphatidylglycerol (PG), (J) Lysophosphatidylcholine (LPC), and (K) Lysophosphatidylethanolamine (LPE). All data are from 3-wk semaglutide treatment. N=6 for both vehicle and semaglutide. ** p<0.01 significant difference vs. Veh.

**Figure S2.**
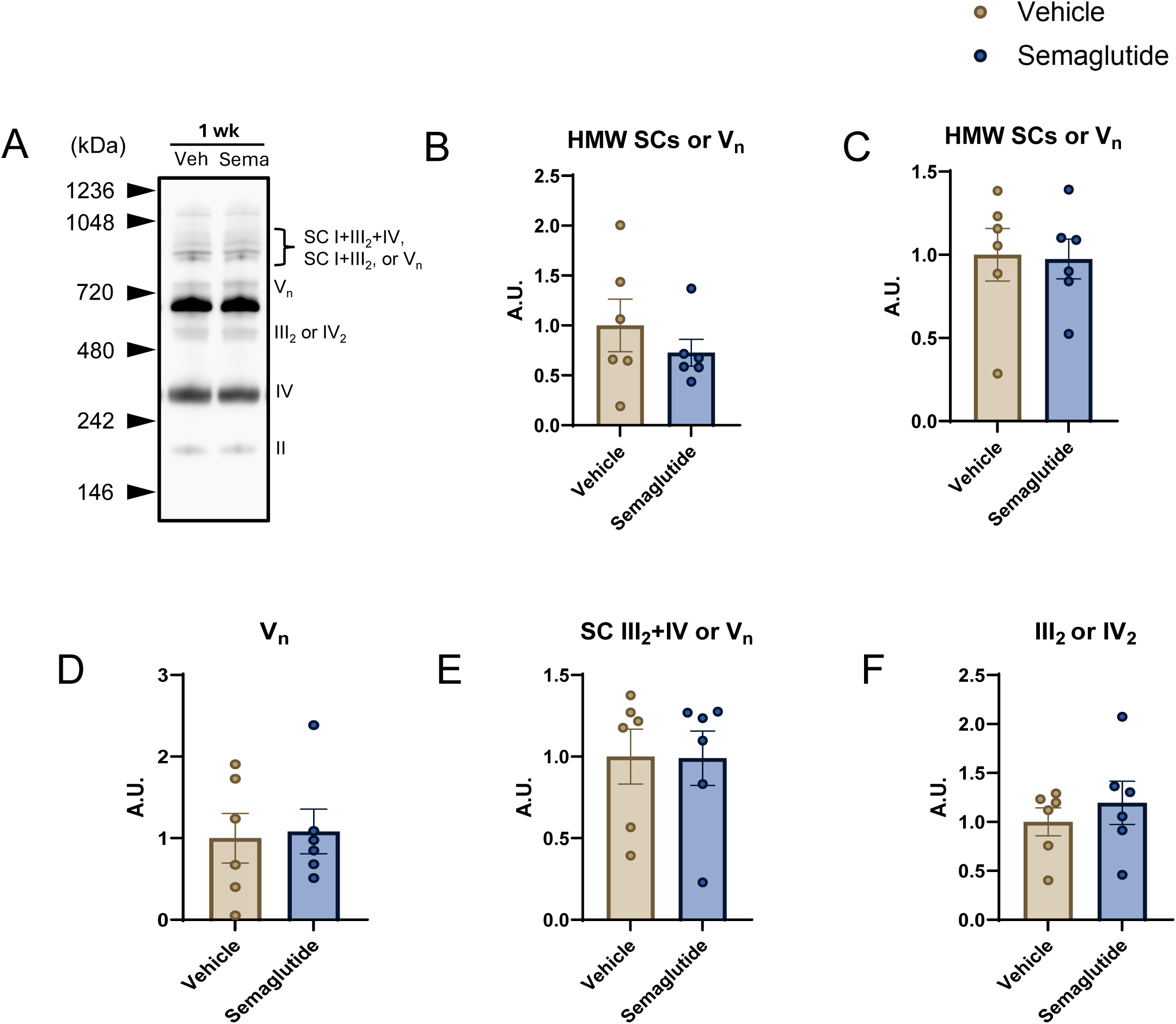
Skeletal muscle mitochondrial supercomplexes. (A) High molecular weight (HMW) supercomplexs (SCs) or V_n_, (B) Supercomplex (SC) I+III_2_+IV and SC I+III_2_ or V_n_,(C) V_n_,(D) SC III_2_+IV or V_n_, (E) III_2_ or IV_2_, (F) IV, and (G) II. All data are from 1-wk semaglutide treatment. N=5-6 for both vehicle and semaglutide.

## References

1. Ryan DH, Yockey SR. Weight Loss and Improvement in Comorbidity: Differences at 5%, 10%, 15%, and Over. Curr Obes Rep. 2017;6(2):187–94. doi: 10.1007/s13679-017-0262-y. PubMed PMID: 28455679; PubMed Central PMCID: PMC5497590.

2. Anderson JW, Konz EC, Frederich RC, Wood CL. Long-term weight-loss maintenance: a meta-analysis of US studies. Am J Clin Nutr. 2001;74(5):579–84. doi: 10.1093/ajcn/74.5.579. PubMed PMID: 11684524.

3. Leibel RL, Rosenbaum M, Hirsch J. Changes in energy expenditure resulting from altered body weight. N Engl J Med. 1995;332(10):621–8. doi: 10.1056/NEJM199503093321001. PubMed PMID: 7632212.

4. Rosenbaum M, Vandenborne K, Goldsmith R, Simoneau JA, Heymsfield S, Joanisse DR, et al. Effects of experimental weight perturbation on skeletal muscle work efficiency in human subjects. Am J Physiol Regul Integr Comp Physiol. 2003;285(1):R183–92. Epub 20030227. doi: 10.1152/ajpregu.00474.2002. PubMed PMID: 12609816.

5. Goldsmith R, Joanisse DR, Gallagher D, Pavlovich K, Shamoon E, Leibel RL, et al. Effects of experimental weight perturbation on skeletal muscle work efficiency, fuel utilization, and biochemistry in human subjects. Am J Physiol Regul Integr Comp Physiol. 2010;298(1):R79–88. Epub 20091104. doi: 10.1152/ajpregu.00053.2009. PubMed PMID: 19889869; PubMed Central PMCID: PMC2806213.

6. Fothergill E, Guo J, Howard L, Kerns JC, Knuth ND, Brychta R, et al. Persistent metabolic adaptation 6 years after “The Biggest Loser” competition. Obesity (Silver Spring). 2016;24(8):1612–9. Epub 20160502. doi: 10.1002/oby.21538. PubMed PMID: 27136388; PubMed Central PMCID: PMC4989512.

7. Baldwin KM, Joanisse DR, Haddad F, Goldsmith RL, Gallagher D, Pavlovich KH, et al. Effects of weight loss and leptin on skeletal muscle in human subjects. Am J Physiol Regul Integr Comp Physiol. 2011;301(5):R1259–66. Epub 20110914. doi: 10.1152/ajpregu.00397.2011. PubMed PMID: 21917907; PubMed Central PMCID: PMC3213951.

8. Joyner MJ, Coyle EF. Endurance exercise performance: the physiology of champions. J Physiol. 2008;586(1):35–44. Epub 20070927. doi: 10.1113/jphysiol.2007.143834. PubMed PMID: 17901124; PubMed Central PMCID: PMC2375555.

9. Murray AJ, Horscroft JA. Mitochondrial function at extreme high altitude. J Physiol. 2016;594(5):1137–49. Epub 20150626. doi: 10.1113/JP270079. PubMed PMID: 26033622; PubMed Central PMCID: PMC4771793.

10. Ferrara PJ, Lang MJ, Johnson JM, Watanabe S, McLaughlin KL, Alan Maschek J, et al. Weight loss increases skeletal muscle mitochondrial energy efficiency in obese mice. Life Metabolism. 2023;2(2). doi: 10.1093/lifemeta/load014.

11. Wilding JPH, Batterham RL, Calanna S, Davies M, Van Gaal LF, Lingvay I, et al. Once-Weekly Semaglutide in Adults with Overweight or Obesity. N Engl J Med. 2021;384(11):989–1002. Epub 20210210. doi: 10.1056/NEJMoa2032183. PubMed PMID: 33567185.

12. Garvey WT, Batterham RL, Bhatta M, Buscemi S, Christensen LN, Frias JP, et al. Two-year effects of semaglutide in adults with overweight or obesity: the STEP 5 trial. Nat Med. 2022;28(10):2083–91. Epub 20221010. doi: 10.1038/s41591-022-02026-4. PubMed PMID: 36216945; PubMed Central PMCID: PMC9556320.

13. Wilding JPH, Batterham RL, Davies M, Van Gaal LF, Kandler K, Konakli K, et al. Weight regain and cardiometabolic effects after withdrawal of semaglutide: The STEP 1 trial extension. Diabetes Obes Metab. 2022;24(8):1553–64. Epub 20220519. doi: 10.1111/dom.14725. PubMed PMID: 35441470; PubMed Central PMCID: PMC9542252.

14. Toledo FG, Menshikova EV, Ritov VB, Azuma K, Radikova Z, DeLany J, et al. Effects of physical activity and weight loss on skeletal muscle mitochondria and relationship with glucose control in type 2 diabetes. Diabetes. 2007;56(8):2142–7. Epub 20070529. doi: 10.2337/db07-0141. PubMed PMID: 17536063.

15. Wang D, Jiang DM, Yu RR, Zhang LL, Liu YZ, Chen JX, et al. The Effect of Aerobic Exercise on the Oxidative Capacity of Skeletal Muscle Mitochondria in Mice with Impaired Glucose Tolerance. J Diabetes Res. 2022;2022:3780156. Epub 20220607. doi: 10.1155/2022/3780156. PubMed PMID: 35712028; PubMed Central PMCID: PMC9197611.

16. Heden TD, Johnson JM, Ferrara PJ, Eshima H, Verkerke ARP, Wentzler EJ, et al. Mitochondrial PE potentiates respiratory enzymes to amplify skeletal muscle aerobic capacity. Sci Adv. 2019;5(9):eaax8352. Epub 20190911. doi: 10.1126/sciadv.aax8352. PubMed PMID: 31535029; PubMed Central PMCID: PMC6739096.

17. Lark DS, Torres MJ, Lin CT, Ryan TE, Anderson EJ, Neufer PD. Direct real-time quantification of mitochondrial oxidative phosphorylation efficiency in permeabilized skeletal muscle myofibers. Am J Physiol Cell Physiol. 2016;311(2):C239–45. Epub 20160622. doi: 10.1152/ajpcell.00124.2016. PubMed PMID: 27335172; PubMed Central PMCID: PMC5129772.

18. Jha P, Wang X, Auwerx J. Analysis of Mitochondrial Respiratory Chain Supercomplexes Using Blue Native Polyacrylamide Gel Electrophoresis (BN-PAGE). Curr Protoc Mouse Biol. 2016;6(1):1–14. Epub 20160301. doi: 10.1002/9780470942390.mo150182. PubMed PMID: 26928661; PubMed Central PMCID: PMC4823378.

19. Bianchi C, Genova ML, Parenti Castelli G, Lenaz G. The mitochondrial respiratory chain is partially organized in a supercomplex assembly: kinetic evidence using flux control analysis. J Biol Chem. 2004;279(35):36562–9. Epub 20040617. doi: 10.1074/jbc.M405135200. PubMed PMID: 15205457.

20. Greggio C, Jha P, Kulkarni SS, Lagarrigue S, Broskey NT, Boutant M, et al. Enhanced Respiratory Chain Supercomplex Formation in Response to Exercise in Human Skeletal Muscle. Cell Metab. 2017;25(2):301–11. Epub 20161201. doi: 10.1016/j.cmet.2016.11.004. PubMed PMID: 27916530.

21. Frey TG, Mannella CA. The internal structure of mitochondria. Trends Biochem Sci. 2000;25(7):319–24. doi: 10.1016/s0968-0004(00)01609-1. PubMed PMID: 10871882.

22. Siripoksup P, Cao G, Cluntun AA, Maschek JA, Pearce Q, Brothwell MJ, et al. Sedentary behavior in mice induces metabolic inflexibility by suppressing skeletal muscle pyruvate metabolism. J Clin Invest. 2024;134(11). Epub 20240423. doi: 10.1172/JCI167371. PubMed PMID: 38652544; PubMed Central PMCID: PMC11142742.

23. Sparks LM, Redman LM, Conley KE, Harper ME, Yi F, Hodges A, et al. Effects of 12 Months of Caloric Restriction on Muscle Mitochondrial Function in Healthy Individuals. J Clin Endocrinol Metab. 2017;102(1):111–21. doi: 10.1210/jc.2016-3211. PubMed PMID: 27778643; PubMed Central PMCID: PMC5413108.

24. Carlessi R, Chen Y, Rowlands J, Cruzat VF, Keane KN, Egan L, et al. GLP-1 receptor signalling promotes beta-cell glucose metabolism via mTOR-dependent HIF-1alpha activation. Sci Rep. 2017;7(1):2661. Epub 20170601. doi: 10.1038/s41598-017-02838-2. PubMed PMID: 28572610; PubMed Central PMCID: PMC5454020.

25. Tsuboi T, da Silva Xavier G, Holz GG, Jouaville LS, Thomas AP, Rutter GA. Glucagon-like peptide-1 mobilizes intracellular Ca2+ and stimulates mitochondrial ATP synthesis in pancreatic MIN6 beta-cells. Biochem J. 2003;369(Pt 2):287–99. doi: 10.1042/BJ20021288. PubMed PMID: 12410638; PubMed Central PMCID: PMC1223096.

26. Kuznetsov AV, Veksler V, Gellerich FN, Saks V, Margreiter R, Kunz WS. Analysis of mitochondrial function in situ in permeabilized muscle fibers, tissues and cells. Nat Protoc. 2008;3(6):965–76. doi: 10.1038/nprot.2008.61. PubMed PMID: 18536644.

27. Picard M, Taivassalo T, Gouspillou G, Hepple RT. Mitochondria: isolation, structure and function. J Physiol. 2011;589(Pt 18):4413-21. Epub 20110627. doi: 10.1113/jphysiol.2011.212712. PubMed PMID: 21708903; PubMed Central PMCID: PMC3208215.

28. Saks VA, Veksler VI, Kuznetsov AV, Kay L, Sikk P, Tiivel T, et al. Permeabilized cell and skinned fiber techniques in studies of mitochondrial function in vivo. Mol Cell Biochem. 1998;184(1-2):81–100. PubMed PMID: 9746314.

29. Kay L, Nicolay K, Wieringa B, Saks V, Wallimann T. Direct evidence for the control of mitochondrial respiration by mitochondrial creatine kinase in oxidative muscle cells in situ. J Biol Chem. 2000;275(10):6937–44. doi: 10.1074/jbc.275.10.6937. PubMed PMID: 10702255.

30. Perry CG, Kane DA, Lin CT, Kozy R, Cathey BL, Lark DS, et al. Inhibiting myosin-ATPase reveals a dynamic range of mitochondrial respiratory control in skeletal muscle. Biochem J. 2011;437(2):215–22. doi: 10.1042/BJ20110366. PubMed PMID: 21554250; PubMed Central PMCID: PMC3863643.

31. He ZH, Bottinelli R, Pellegrino MA, Ferenczi MA, Reggiani C. ATP consumption and efficiency of human single muscle fibers with different myosin isoform composition. Biophys J. 2000;79(2):945–61. doi: 10.1016/S0006-3495(00)76349-1. PubMed PMID: 10920025; PubMed Central PMCID: PMC1300991.

32. Edman S, Flockhart M, Larsen FJ, Apro W. Need for speed: Human fast-twitch mitochondria favor power over efficiency. Mol Metab. 2024;79:101854. Epub 20231215. doi: 10.1016/j.molmet.2023.101854. PubMed PMID: 38104652; PubMed Central PMCID: PMC10788296.

33. Luna-Marco C, de Maranon AM, Hermo-Argibay A, Rodriguez-Hernandez Y, Hermenejildo J, Fernandez-Reyes M, et al. Effects of GLP-1 receptor agonists on mitochondrial function, inflammatory markers and leukocyte-endothelium interactions in type 2 diabetes. Redox Biol. 2023;66:102849. Epub 20230814. doi: 10.1016/j.redox.2023.102849. PubMed PMID: 37591012; PubMed Central PMCID: PMC10457591.

34. Jastroch M, Divakaruni AS, Mookerjee S, Treberg JR, Brand MD. Mitochondrial proton and electron leaks. Essays Biochem. 2010;47:53-67. doi: 10.1042/bse0470053. PubMed PMID: 20533900; PubMed Central PMCID: PMC3122475.

35. Lin TK, Lin KJ, Lin HY, Lin KL, Lan MY, Wang PW, et al. Glucagon-Like Peptide-1 Receptor Agonist Ameliorates 1-Methyl-4-Phenyl-1,2,3,6-Tetrahydropyridine (MPTP) Neurotoxicity Through Enhancing Mitophagy Flux and Reducing alpha-Synuclein and Oxidative Stress. Front Mol Neurosci. 2021;14:697440. Epub 20210707. doi: 10.3389/fnmol.2021.697440. PubMed PMID: 34305527; PubMed Central PMCID: PMC8292641.

36. Nunn E, Jaiswal N, Gavin M, Uehara K, Stefkovich M, Drareni K, et al. Antibody blockade of activin type II receptors preserves skeletal muscle mass and enhances fat loss during GLP-1 receptor agonism. Mol Metab. 2024;80:101880. Epub 20240111. doi: 10.1016/j.molmet.2024.101880. PubMed PMID: 38218536; PubMed Central PMCID: PMC10832506.

37. Zurlo F, Larson K, Bogardus C, Ravussin E. Skeletal muscle metabolism is a major determinant of resting energy expenditure. J Clin Invest. 1990;86(5):1423–7. doi: 10.1172/JCI114857. PubMed PMID: 2243122; PubMed Central PMCID: PMC296885.

38. Sanchez E, Lobo T, Fox JL, Zeviani M, Winge DR, Fernandez-Vizarra E. LYRM7/MZM1L is a UQCRFS1 chaperone involved in the last steps of mitochondrial Complex III assembly in human cells. Biochim Biophys Acta. 2013;1827(3):285–93. Epub 20121117. doi: 10.1016/j.bbabio.2012.11.003. PubMed PMID: 23168492; PubMed Central PMCID: PMC3570683.

39. Bottani E, Cerutti R, Harbour ME, Ravaglia S, Dogan SA, Giordano C, et al. TTC19 Plays a Husbandry Role on UQCRFS1 Turnover in the Biogenesis of Mitochondrial Respiratory Complex III. Mol Cell. 2017;67(1):96–105 e4. Epub 20170629. doi: 10.1016/j.molcel.2017.06.001. PubMed PMID: 28673544.

40. Cherian A, Divya KP, Jose J, Thomas B. Multifocal cavitating leukodystrophy-A distinct image in mitochondrial LYRM7 mutations. Mult Scler Relat Disord. 2021;47:102615. Epub 20201105. doi: 10.1016/j.msard.2020.102615. PubMed PMID: 33189022.

41. Ghezzi D, Arzuffi P, Zordan M, Da Re C, Lamperti C, Benna C, et al. Mutations in TTC19 cause mitochondrial complex III deficiency and neurological impairment in humans and flies. Nat Genet. 2011;43(3):259–63. Epub 20110130. doi: 10.1038/ng.761. PubMed PMID: 21278747.

42. Invernizzi F, Tigano M, Dallabona C, Donnini C, Ferrero I, Cremonte M, et al. A homozygous mutation in LYRM7/MZM1L associated with early onset encephalopathy, lactic acidosis, and severe reduction of mitochondrial complex III activity. Hum Mutat. 2013;34(12):1619–22. Epub 20130923. doi: 10.1002/humu.22441. PubMed PMID: 24014394; PubMed Central PMCID: PMC4028993.

